# Orbitofrontal cortex and learning predictions of state transitions

**DOI:** 10.1101/2020.09.17.302521

**Authors:** Stephanie C.Y. Chan, Nicolas W. Schuck, Nina Lopatina, Geoffrey Schoenbaum, Yael Niv

## Abstract

Learning the transition structure of the environment – the probabilities of transitioning from one environmental state to another – is a key prerequisite for goal-directed planning and model-based decision making. To investigate the role of the orbitofrontal cortex (OFC) in goal-directed planning and decision making, we used fMRI to assess univariate and multivariate activity in the OFC while humans experienced state transitions that varied in degree of surprise. Converging with recent evidence, we found that OFC activity was related to greater learning about transition structure. However, the observed relationship was inconsistent with a straightforward interpretation of OFC activity as representing a state prediction error that would facilitate learning of transitions via error-correcting mechanisms. The state prediction error hypothesis predicts that OFC activity at the time of observing an outcome should increase expectation of that same observed outcome on subsequent trials. Instead, our results showed that OFC activity was associated with increased expectation of the *more probable* outcome; that is, with more optimal predictions. These results suggest that the OFC is involved in updating or reinforcing a learned transition model on a trial-by-trial basis, specifically for the currently observed cue-outcome associations. Our findings add to the evidence of OFC involvement in learning state-to-state transition structure, while providing new constraints for algorithmic hypotheses regarding how these transitions are learned.

**Significance Statement:** The orbitofrontal cortex (OFC) has been implicated in model-based decision making—the kind of decisions that result from planning using an “environment model” of how current actions affect our future states. However, the widely suggested role of the OFC in representing expected values of future states is not sufficient to explain why the OFC would be critical for planning in particular. A new line of evidence implicates the OFC in learning about transition structure of the environment – a key component of the “environment model” used for planning. We investigate this function, adding to the growing literature on the role of the OFC in learning and decision making, while unveiling new questions about the algorithmic role of OFC in goal-directed planning.

## Introduction

To flexibly plan for the future, we must be able to predict which states of the world lead to which (i.e. we need to learn a model of the “transition structure” of the world). For example, to decide whether to drink warm milk or coffee, we need to know that warm milk makes us sleepy, but coffee wakes us up. This type of planning has been termed “model-based decision making”, in contrast to “model-free decision making”, which does not require such a model (Daw et al, 2005).

The orbitofrontal cortex (OFC) has been shown to be particularly important for model-based decision-making (Baxter et al, 2000; Izquierdo et al, 2004; Valentin et al, 2007; De Wit et al, 2009; Walton et al, 2010; McDannald et al, 2011; Rudebeck et al, 2011). However, previous research has focused on showing that OFC activity relates to the expected values of future rewards (Gottfried et al, 2003; Padoa-Schioppa and Assad, 2006; Hampton et al, 2006; Fellows, 2007; Hare et al, 2008; Wallis and Kennerley, 2011; Monosov and Hikosaka, 2012). Recently, we have instead proposed that the OFC represents the current state of the task (Schuck et al, 2016), and that the OFC is especially critical for making decisions in situations where environmental stimuli do not unambiguously determine the task-relevant state (e.g., whether the state is “Thursday evening” and it is bedtime, versus “Friday evening,” in which case I don’t want to become sleepy as I am going to a party; Wilson et al, 2014; Bradfield et al, 2015; Chan et al., 2016; Nogueira et al, 2017). However, both value and state representation are important in model-free as well as model-based decision making, and therefore these two lines of research do not explain why the OFC is critical specifically for the latter.

Yet another line of research provides a potential explanation for the OFC’s particular prominence in model-based planning. This research suggests that the OFC is important for learning about the state-to-state “transition structure” of the world – the tendencies of certain environmental states to lead to other states. One study showed that OFC-lesioned rats couldn’t learn about changes in the transitions from cues to outcomes (cue-outcome associations; McDannald et al, 2011), while a study in humans linked fMRI surprise signals in lateral OFC with updates in hippocampus of a model of transition structure (Boorman et al, 2016). Some newer studies have observed such surprise signals in the midbrain (Sharpe et al, 2017; Takahashi et al, 2017; Stalnaker et al, 2019), or in a set of areas including the midbrain and lateral OFC (Howard and Kahnt, 2018; Suarez et al, 2019). Howard and Kahnt (2018) additionally found that midbrain identity prediction errors were correlated with cue-outcome learning and changes in outcome identity representations in the OFC. The hypothesized link between OFC and learning transition structure could also explain OFC’s centrality to model-based decision making, given that transition structure is a critical component of the “model” in such decision making. One cannot plan and mentally simulate the future result of current actions without an accurate model of how state transitions are likely to unfold in the future.

How exactly might the OFC be involved in learning about transition structure? The OFC might itself compute or represent a prediction error at the time of unexpected outcomes, which can be used to update an internal model of transition structure. Such “state prediction error” signals would occur upon observing state transitions that are unexpected, and could be used to guide learning so that transitions are better predicted in the future (e.g. Glascher et al, 2010). Note that these error signals are analogous to – but distinct from – reward prediction errors that are used for learning to associate states with their reward values (e.g., Rescorla and Wagner, 1972; Montague et al, 1996). However, the existing research does not make specific predictions about the role of OFC in representing or learning about transition structure, and state prediction errors are just one possible way. We therefore set out here to test whether the OFC might be in involved in error-driven learning via signaling of state prediction errors, and whether OFC activity could predict behavior related to learning transition structure. We also tested the two dominant hypotheses of OFC function – representing the current state and representing expected value.

In our experiment, black-and-white image cues led stochastically to M&M candies of different quantities and colors (outcomes). In the critical trials, the number of M&Ms was fully predictable, but their color was not, so as to generate state prediction errors in the absence of reward prediction errors. Using fMRI, we investigated activity in the human OFC at the time of these outcomes, and its relationship with participants’ behavioral predictions of state transitions.

## 2 Materials and Methods

### 2.1 Subjects

Twenty-four volunteers from the Princeton University community participated in exchange for monetary compensation ($20 per hour + up to $10 performance-related bonus). All subjects were right-handed (14 female, age range 18-34 years) and stated that they liked M&Ms. Informed written consent was obtained from all subjects, and the study protocol was approved by the Institutional Review Board for Human Subjects at Princeton University.

### 2.2 Experimental design

Each trial began with 0.5 - 8 seconds of fixation (truncated exponential distribution, mean 2.4 s). Then one of four black-and-white image cues depicting outdoor scenes appeared for 1.2 s (see Fig 1a). On 75% of the trials, this was followed by the opening of a box around the image (0.2 s). Then, a set of M&Ms appeared below the image and fell into a bowl, over the course of 0.9 s. As the M&Ms fell into the bowl, one clinking sound was emitted for each M&M in the set. A tally at the bottom of the screen (not shown in Fig 1a) indicated the total number of M&Ms received so far, for each of the four possible colors.

**Figure 1.**
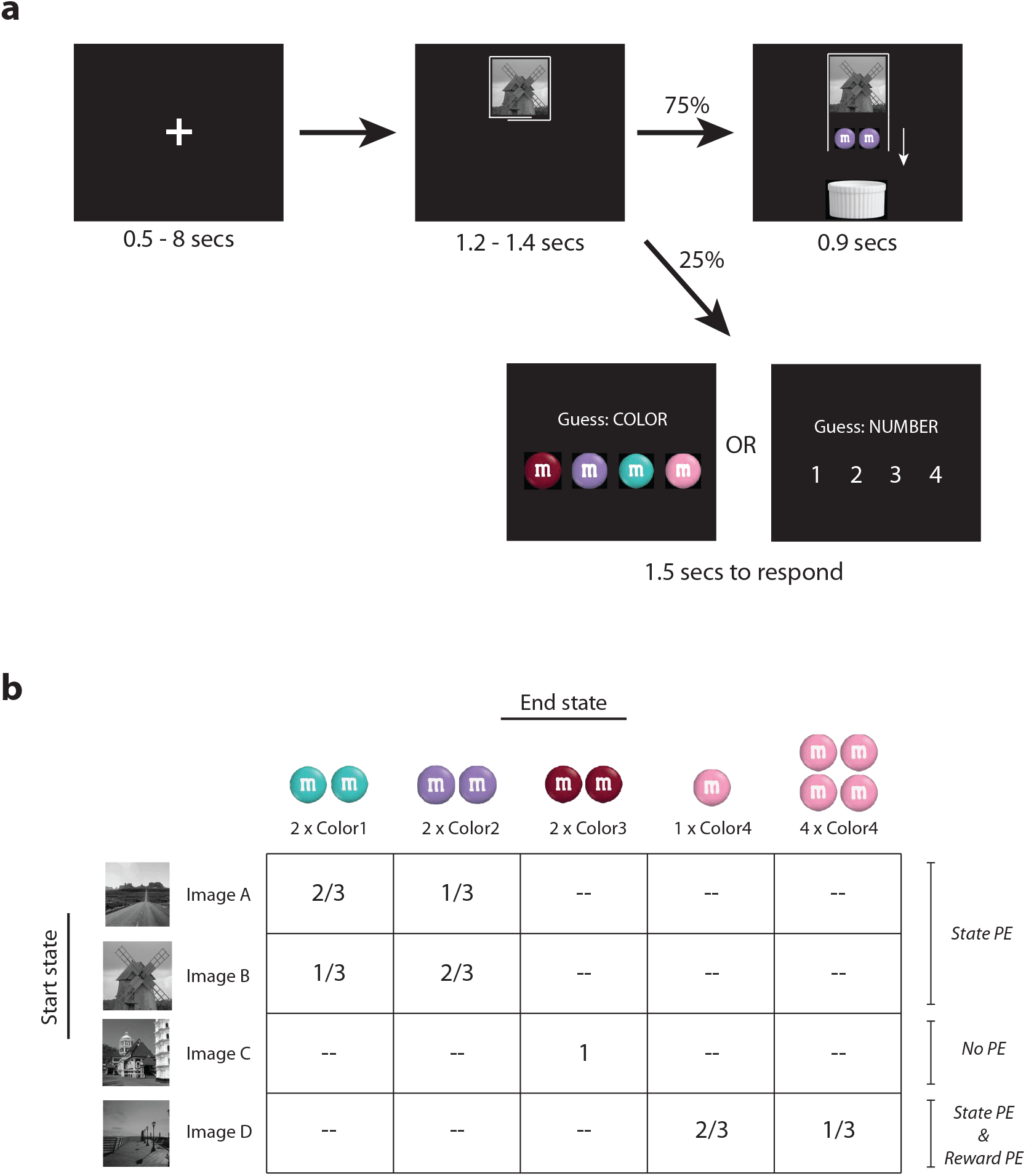
**(a) Experimental design.** Trials began with fixation. Then, one of four image cues (4 different “start states”) appeared. On most trials, the box around the image opened, and a number of colored M&Ms dropped from the image, clinking as they fell into a bowl (5 different “end states”). On the randomly interspersed “guess” trials, the image cue was instead followed by a prompt to guess (within 1.5 seconds) either the color or number of M&Ms that would have fallen on that trial. **(b) Cue-outcome contingencies for each of the four images (transition matrix for the experiment)**. Numbers in table indicate probability of each end state (M&M outcome) given each start state (image cue). PE = prediction error. Larger state prediction errors are expected for rarer outcomes (smaller transition probabilities). Images and M&M colors were assigned randomly for each subject. Our analyses focused on Cue A and Cue B trials, which were designed to elicit state prediction errors in the absence of reward prediction errors.

Each of the four image cues was associated with different numbers and colors of M&Ms according to a predetermined schedule of reinforcement (Fig 1b). Cue A and Cue B were designed to elicit state prediction errors throughout the experiment due to a probabilistic schedule of M&M color, but not reward prediction errors, because they always dropped exactly 2 M&Ms. Cue C, in contrast, was associated with 2 M&Ms of a fixed color, thus eliciting no prediction errors once the contingencies had been learned. Finally, Cue D was designed to elicit only reward prediction errors—it dropped either 1 or 4 M&Ms of a fixed color (as with the other image cues, Cue D led to 2 M&Ms on average, such that all 4 cues were equated for average reward value). For each subject, the images and M&M colors were assigned randomly from a pool of 20 images and 5 M&M colors.

Subjects earned one real M&M of a given color for every 17 “virtual” M&Ms that they received in the task. Subjects were requested to refrain from eating or drinking (except water) for at least 3 hours prior to the experiment, so that the M&Ms would be especially rewarding. Non-standard M&M colors were chosen to circumvent pre-existing preferences for specific M&M colors, and to achieve perceptually distinct outcomes that are of equal value. Note also that our analyses of state prediction error always combine Cue A and Cue B trials, so that any potential value differences between the two colors cancel out. In a post-experiment questionnaire, subjects rated the appeal of the M&Ms on a scale from 1 (not appealing at all) to 5 (very appealing). The mean rating was 3.8 ± 0.2.

25% of all trials (pseudorandomly distributed) were “guess trials”. On these trials, the appearance of the black-and-white image cue was followed by a prompt reading “Guess: COLOR” or “Guess: NUMBER”. At the appearance of the prompt, the image cue disappeared. Subjects were given 1.5 s to guess what color/number of M&Ms *would have* fallen on that trial. Subjects received 10¢ for every question correctly answered. The purpose of the guess trials was to encourage subjects to pay attention to the image cue and to actively make a prediction of the upcoming M&M outcome on *every* trial – because the allowed response time was so short, subjects had to prepare an answer upon viewing the image cue in case a guess prompt followed.

Subjects performed 72 training trials outside of the scanner, to familiarize themselves with the task and to learn the stimulus-outcome contingencies. During training, subjects received and ate the M&Ms they earned (approximately 7 M&M candies). They were then informed that future M&Ms they earned would be given to them after the ensuing scanning session, and they performed another 420 trials in the MRI scanner. At the end of the experiment, subjects received all M&Ms earned while in the scanner. The 420 trials were evenly distributed between the four image cues, with trial order pseudorandomized so that the total number of M&Ms collected increased at the same rate for every color. The experiment was divided into 5 scan sessions of approximately 10 minutes each.

### 2.3 Behavioral measures

We evaluated three types of behavioral measures, computed separately for each subject and for each prediction trial type (image cue type x number/color prediction): (1) overall performance over the course of the experiment; (2) change in performance over the course of the experiment (3) sensitivity to the most recent outcome (a proxy for learning rate).

To assess overall performance, we computed the fraction of responses that were optimal (i.e. for which the subject selected the common outcome), across all scan sessions. To measure change in performance, we computed the difference in performance from the beginning to the end of the experiment as the fraction of optimal responses in the last scan session minus the fraction of optimal responses in the training session. To assess sensitivity to previous outcome, we computed the probability of predicting the common outcome after observing the common outcome on the previous trial with the same image cue, compared to the probability of predicting the common outcome after observing the uncommon outcome on the previous trial with the same image cue. The difference between these two quantities served as a proxy for learning rate – subjects with high learning rate would be more sensitive to the most recent outcome and would show a larger difference between the two quantities.

### 2.4 fMRI acquisition

Functional brain images were acquired using a 3T MRI scanner (Skyra; Siemens Erlangen, Germany), and were preprocessed using FSL (http://fsl.fmrib.ox.ac.uk/fsl/). An echoplanar imaging sequence was used to acquire 40 slices of 2mm thickness with a 1-mm gap (repetition time (TR) = 2.4s, echo time (TE) = 27ms, flip angle = 71°, field of view = 196 mm, phase encoding direction = anterior to posterior). We optimized our fMRI sequence for OFC signal acquisition by including a gap between slices, using shimming and fieldmap unwarping, and tilting the slices by approximately 30° from the axial plane towards a coronal orientation (Deichmann et al, 2003). Fieldmaps consisted of forty 3-mm slices, centered at the centers of the echoplanar slices, with TR = 500ms, TE1 = 3.99 ms, TE2 = 6.45ms, field of view = 196mm. At the end of the 5 functional scanning sessions, an MPRAGE anatomical scan was acquired, consisting of 176 1-mm axial slices, TR = 2.3s, TE = 3.08 ms, flip angle = 9°, and field of view = 256mm.

### 2.5 Preprocessing

All functional images were preprocessed using high pass filtering (filter at 1/100 Hz), motion correction (six-parameter rigid body transformation), correction for B0 magnetic inhomogeneities (fieldmap unwarping), spatial smoothing (Gaussian kernel with full width at half maximum of 5mm), and co-registration of functional and structural scans. For GLM results, we additionally performed spatial normalization of subject-level results to match a template in MNI space (12-parameter affine transformation).

### 2.6 Functional parcellation of orbitofrontal cortex

Regions of interest for the orbitofrontal cortex were obtained from Kahnt et al. (2012), who used k-means clustering of functional connectivity patterns to parcellate OFC into subregions. We used the parcellation of OFC into two clusters, which correspond with medial-lateral subdivisions of OFC found in studies of cytoarchitectonic structure and of intra-regional anatomical connectivity (Carmichael and Price, 1996; Ongür and Price, 2000).

### 2.7 Obtaining mean percent signal change at M&M outcomes

Using the FSL toolbox (http://fsl.fmrib.ox.ac.uk/fsl/), we performed a GLM analysis with the following regressors: one regressor for the onsets of each type of image cue (A, B, C, D); one regressor for the onsets of the M&M outcomes for Cue C; one regressor for the onsets of the uncommon outcomes for each of the image cues A, B, and D (3 regressors total); one regressor for the onsets of the common outcomes for each of the image cues A, B, and D (3 regressors total); and one parametric regressor for the clinks of the M&Ms into the bowl (1, 2, or 4 clinks). These 12 regressors were convolved with a standard hemodynamic response function. In addition, the design matrix included 6 motion regressors and an intercept (constant) term.

Regressor weights for each voxel and each scan session were converted to percent signal change by multiplying by the appropriate scale factor for events of length 0.1 sec convolved with the standard double-gamma hemodynamic response function, and then dividing by the mean of the voxel’s timecourse for that scan session. These per-scan numbers were averaged across scans for each subject. To obtain the percent signal change for a region of interest, the percent signal change was averaged across all voxels in the region of interest.

### 2.8 Obtaining trial-by-trial estimates of percent signal change at M&M outcomes

To obtain trial-by-trial estimates of percent signal change (PSC) in an ROI at each M&M outcome, we fit a separate GLM for each trial. This GLM was identical to the one used for estimating mean PSC (above), except that the regressor for the condition of the trial of interest was split into two – one regressor modeled the onset for the trial of interest only, and a second regressor modeled the onsets of all other trials in that condition (Mumford et al, 2012). These GLMs were fitted to data that were preprocessed in FSL, but the GLMs themselves were fitted using in-lab code written in MATLAB, for computational reasons.

### 2.9 MVPA classification

The purpose of our MVPA analyses was to test whether activity in OFC at the time of the M&M outcomes contained information about the start state and end state (stimulus and outcome) for each transition. We analyzed the trials that were designed to elicit state prediction errors (Cue A and Cue B trials).

Given our rapid event-related design, we first used a GLM to deconvolve neighboring events, regress out motion artifacts, and to de-noise examples through averaging (Mumford et al, 2012). To maximize power, we divided each scan in two (1^st^ and 2^nd^ half). The GLM then included, for each half of each scan session, regressors modeling the appearance of the M&Ms for each of four trial types of interest (Cue A followed by M&M Color 1, Cue A followed by M&M Color 2, Cue B followed by M&M Color 1, Cue B followed by M&M Color 2), totaling 8 regressors per scan. The regressors were convolved with a canonical hemodynamic response function. In addition, for each scan session we modeled head motion using six motion regressors and the mean activity using an intercept regressor. We estimated this GLM on each subject’s smoothed, motion-corrected fMRI data using the FSL toolbox (http://fsl.fmrib.ox.ac.uk/fsl/).

We used the resulting patterns of voxel-wise regressor weights for the four trial types (two regressor weights per run and trial type; z-scored within voxels) as training and testing examples for a support vector machine (SVM) classification algorithm with a linear kernel (nu-SVM, as implemented in LIBSVM; Chang and Lin, 2011), under a leave-one-out cross validation scheme, using the Princeton MVPA Toolbox (https://code.google.com/p/princeton-mvpa-toolbox). We used a standard cost (nu) parameter of 1 for the SVM (results did not depend strongly on this parameter).

To classify start state, we classified training and testing examples according to the image cue (Cue A or Cue B). To classify end state, we classified training and testing examples according to the M&M color (Color 1 or Color 2).

## 3 Results

### 3.1 Behavioral performance

For the prediction task, the optimal strategy was to predict the most common outcome on every trial, given that this would maximize subjects’ payout. Overall, subjects predicted the most common outcome 77 ± 2% of the time. The 23% non-optimal guesses may have resulted from a combination of probability matching (for probabilistic transitions, Vulkan, 2000; Erev and Barron, 2000), imperfect knowledge of transition probabilities, and noise. Fig 2a shows subjects’ performance on each trial type. Subjects performed significantly above chance for all trial types (p < 10^-6; one-sided bootstrap test).

**Figure 2.**
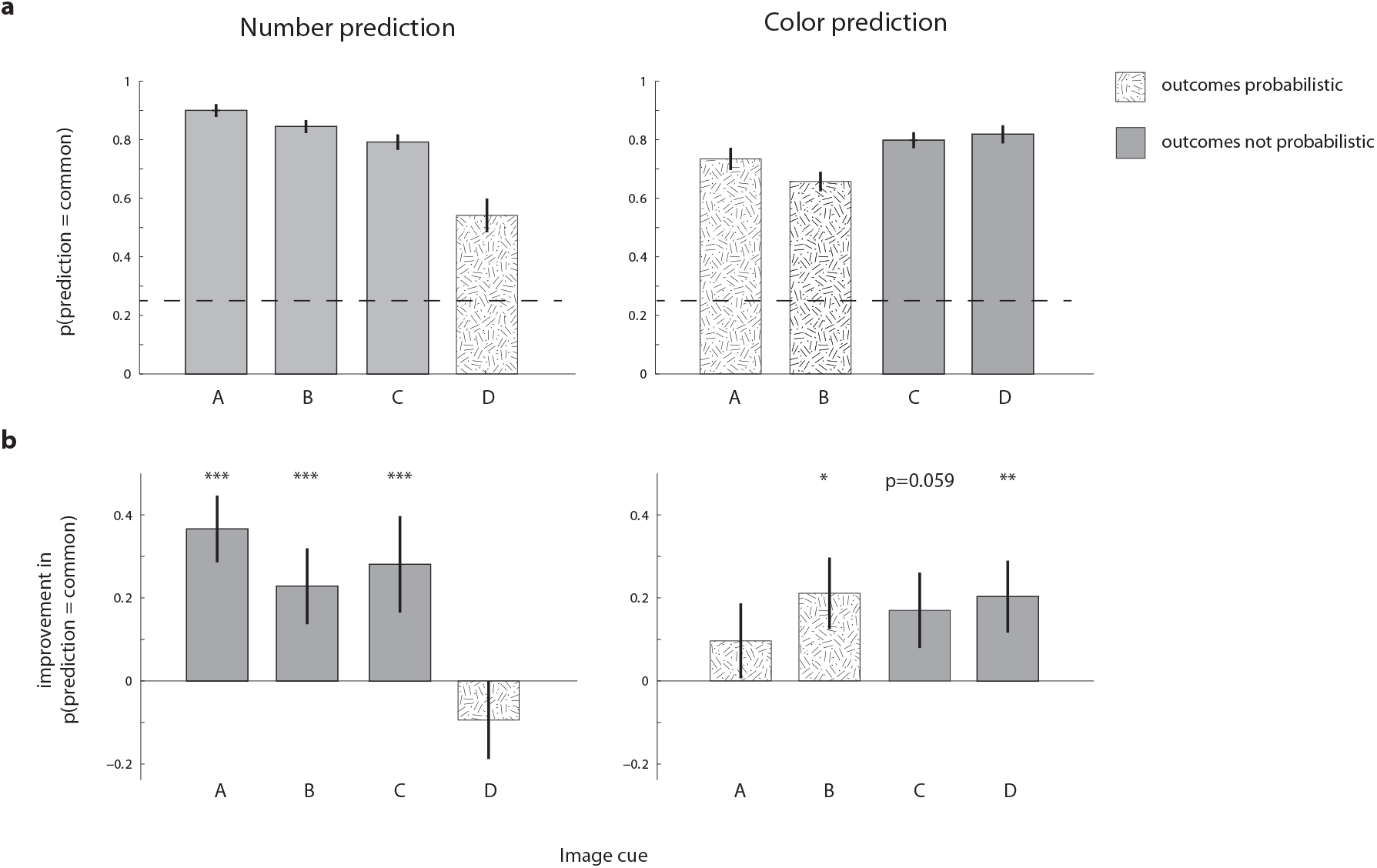
Behavioral performance evidenced learning throughout the task. Hatched bars indicate that the outcomes were probabilistic for that cue and dimension (i.e. Cue D for number, and Cues A and B for color). Error bars indicate standard error of the mean. **(a) Subjects performed above chance in all prediction tasks**. Plotted is the probability of choosing the more common outcome (the optimal prediction) for number prediction trials and color prediction trials across the whole experiment. Dashed line: chance. **(b) Subjects’ performance improved significantly across the experiment, except for color prediction for cue C (which only improved numerically) and number prediction for Cue D**. Plotted is the difference in probability of choosing the more common outcome in the last session compared to the training session. Positive differences indicate learning during the task. *p < 0.05, **p < 0.01 ***p < 0.0001

### 3.2 Overall learning across the experiment

Subjects predicted the more common outcome more often as the experiment progressed, as measured by the difference between performance on the last scan session compared to performance during the training session (before entering the scanner) (Fig 2b). The only exception was in predicting the number of M&Ms for Cue D. Here, the optimal prediction was 1 M&M; however, participants predicted this amount on only around half the prediction trials and predicted the rare 4 M&Ms otherwise, possibly because of the high salience and appeal of the 4 M&Ms outcome. That is, although the 4 M&M outcome was delivered on only 1/3 of the trials involving Cue D, participants may have been confused regarding its frequency, or they may have predicted 4 M&Ms as a form of “wishful thinking”. Over the course of the task, predictions of the outcome of this cue did not improve, and even got worse numerically (Fig 2b).

### 3.3 Learning from recent outcomes

We evaluated each subject’s sensitivity to the most recent outcome as a behavioral proxy for learning rate – a subject with a high learning rate should be relatively more likely to expect an outcome that she recently experienced, while a subject with a low learning rate should be less affected by recent experience. To measure this, we compared the probability of the subject predicting the common outcome for a specific cue after most recently experiencing the common outcome for that cue, versus after most recently experiencing the uncommon outcome. Stronger sensitivity to the most recent outcome, i.e. higher learning rates, should manifest as larger differences between the two quantities. We evaluated learning for the scan sessions, as these were the sessions for which we could correlate learning with brain activity.

For color prediction on Cue A and Cue B trials, subjects showed significantly greater probability of choosing the common outcome if the most recent outcome was common, suggesting that subjects were learning about Cue A and B outcomes from experience during the scan sessions (Fig 3a, left). This pattern of learning was not apparent for Cue D number prediction trials, consistent with the low overall accuracy and low improvement across the experiment for predicting the number of M&Ms for Cue D (Fig 3a, right).

**Figure 3.**
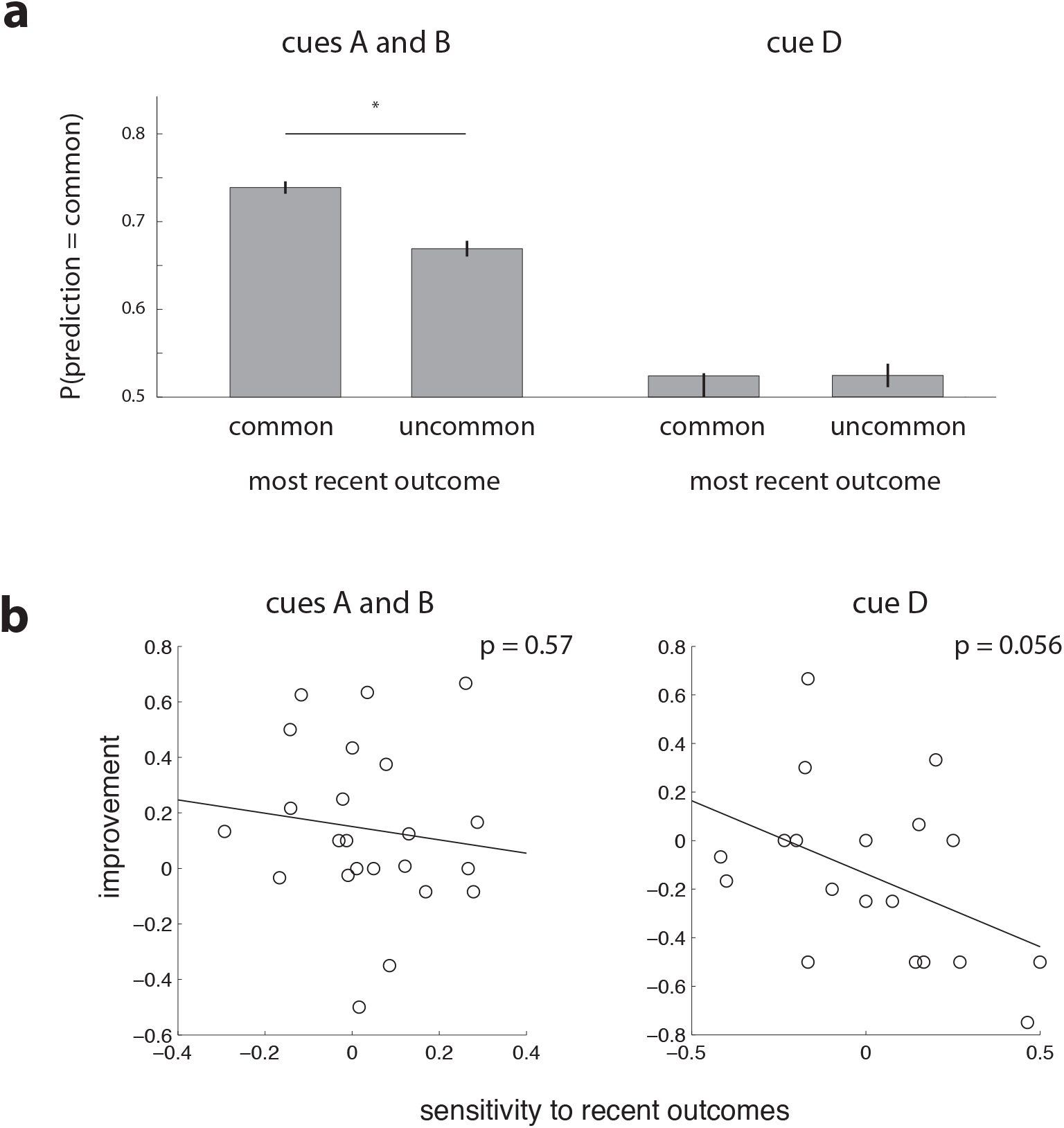
Subjects’ predictions were sensitive to the most recent outcomes for Cues A and B, evidencing trial-by-trial learning. **(a)** Subjects were more likely to predict the common color (for cue A and B trials, where color of the M&M outcome varied; left) after observing the common outcome (as opposed to the uncommon outcome) on the most recent trial with the same image cue. This pattern did not hold for predictions of number in the condition where number of M&Ms varied (number prediction on Cue D trials; right). Means ± SEM. **(b)** There was no correlation between sensitivity to recent outcomes (calculated as the difference between the probability of predicting the common outcome after recently observing the common outcome versus the uncommon outcome for the same cue; see panel a) and performance improvement across the experiment (calculated as the difference in proportion of optimal predictions between the last session and the training session; see Figure 2b).

Note that higher sensitivity to recent outcomes does not necessarily imply greater improvement in performance across the experiment, because high learning rates can, in fact, lead to more highly fluctuating responses. Indeed, as shown in Fig 3b, we did not find evidence for correlation between sensitivity to recent outcomes and improvement across the experiment in Cue A and B color prediction, and we found a marginally negative correlation with improvement in Cue D number prediction.

### 3.4 Identity of outcomes (but not of image cues) was decodable from multivariate OFC activity – OFC does not simply represent perceptual input

To evaluate OFC representations of the current state, we used multivariate classification methods to classify the outcome states (Color 1 vs Color 2) at the time of the M&M outcome for Cue A and Cue B trials. We analyzed a pre-defined OFC region of interest (Figure 4a). Cross-validated classifier performance was significantly above chance (50%) for classifying M&M outcome (classification accuracy 53.9% and 54.0%, p=0.013 and 0.012, for medial and lateral OFC respectively; one-sided bootstrap test), indicating reliable representations of outcome state in both medial and lateral OFC (Fig 4b). In contrast, we did not find above-chance classifier performance for the image cue (Cue A vs Cue B) at the time of the outcome (classification accuracy 49.6% and 48.5%, p=0.61 and 0.75, for medial and lateral OFC respectively; one-sided bootstrap test). This is despite the fact that, on each trial, the image cue was still on the screen at the time that the M&M outcome appeared, and in fact occupied a much larger area of the screen than the M&Ms, indicating that OFC representations of the current state do not simply reflect perceptual input.

**Figure 4.**
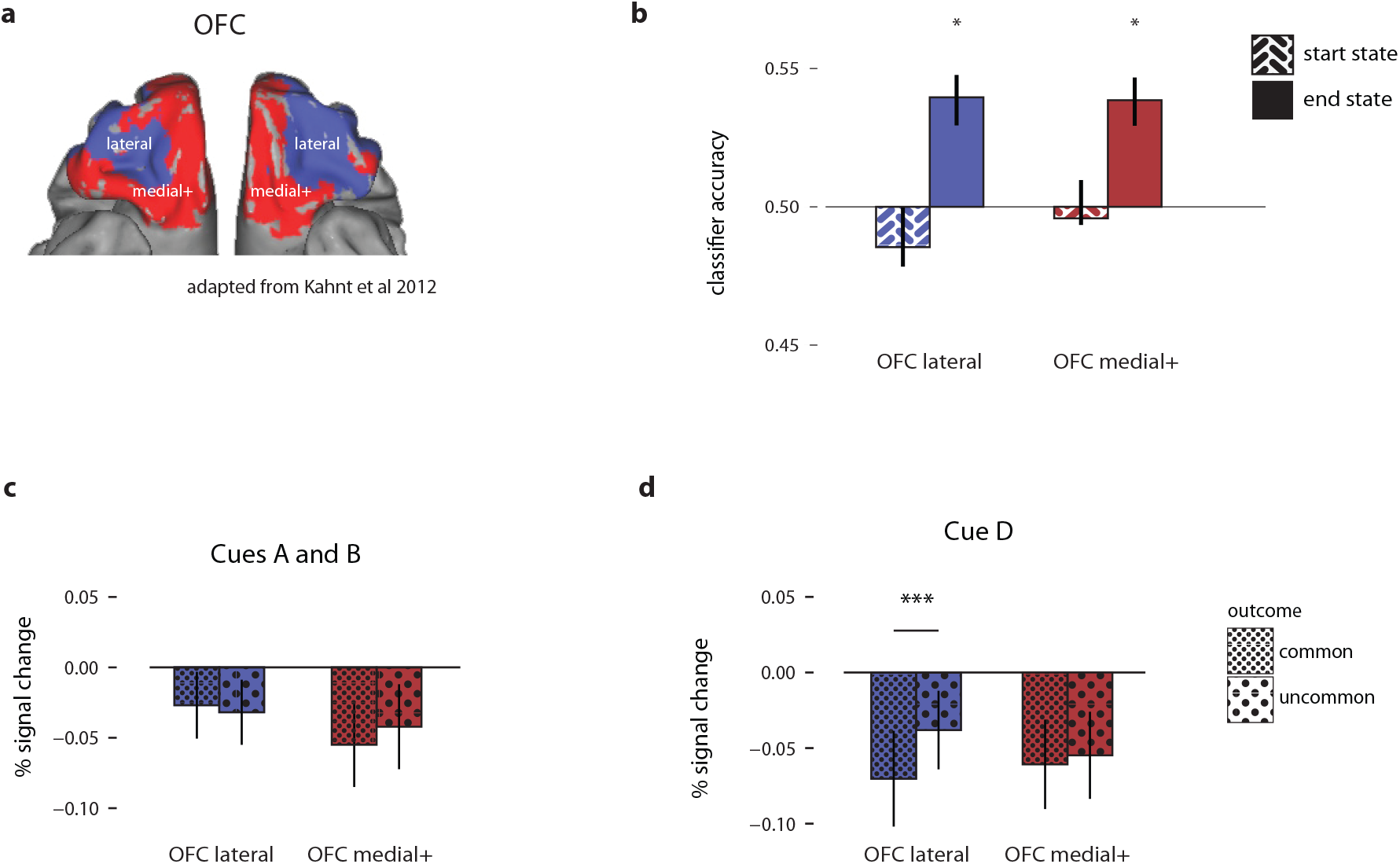
M&M outcomes, but not image cues, were classifiable from multivariate OFC patterns of activity. **(a)** Subregions of OFC, displayed on the orbital surface of the brain. These regions of interest were obtained on a different dataset by Kahnt et al (2012), who parcellated the OFC using k-means clustering of functional connectivity. **(b)** Cross-validated classification performance (across subjects) for start state (image cue) and end state (M&M color) for Cue A and B trials, using multivariate linear classifiers on OFC activity. Error bars indicate SEM. *p < 0.05. **(c-d)** No significant differences in univariate BOLD responses for common vs. uncommon outcomes were observed, except for Cue D (for which uncommon outcomes also corresponded to larger rewards) in lateral OFC. Plotted is percent signal change in subregions of OFC at the time of the common outcomes and the uncommon outcomes. ***p < 0.005

### 3.5 Univariate OFC responses at the time of outcome did not signal state prediction errors

In general, we did not observe significant differences in univariate BOLD responses for common vs. uncommon outcomes (corresponding to hypothesized small vs. large prediction errors). The exception was in lateral OFC for Cue D, where the BOLD response was more negative for the common (1 M&M) outcome as compared to the uncommon 4 M&M outcome (Fig. 4c-d; p<0.005, one-sided bootstrap test), suggesting possible sensitivity to reward value or salience in lateral OFC in particular.

### 3.6 Trial-by-trial correlations of OFC activity with learning from the most recent outcome

Given that subjects’ behavior demonstrated learning from the most recent outcome for Cues A and B during the scan sessions (Fig 3a, section 3.3), we evaluated whether OFC activity could predict this learning, on a trial-by-trial basis. For each subject, we used logistic regression on OFC activity at the time of an outcome to predict whether the subject would choose the common outcome in the subsequent “guess” trial involving the same cue. The results of this analysis indicated an involvement of OFC in learning about transitions, but they were not consistent with a straightforward interpretation of OFC activity as reflecting a state prediction error.

Based on a prediction-error account of OFC, we would expect that the slope term of the logistic regression would be positive for the common outcomes and negative for the uncommon outcomes—that is, if greater OFC activity at the time of an outcome indicates a larger prediction error (and therefore more learning to predict that outcome), then it should lead to a greater probability of the subject predicting the *same* outcome on the next trial (common after observing a common outcome, and uncommon after observing an uncommon outcome). Instead, we found that the fit slope terms were positive for both trials where the most recent outcome was the common outcome, as well as for trials where the most recent outcome was the uncommon outcome. In other words, no matter the outcome (common or uncommon), greater BOLD activity in OFC at the time of an outcome was correlated with higher probability of subjects predicting the *common* outcome on the next trial with the same cue (Fig 5a). These results suggest the OFC’s involvement on a trial-by-trial basis in learning task contingencies.

**Figure 5.**
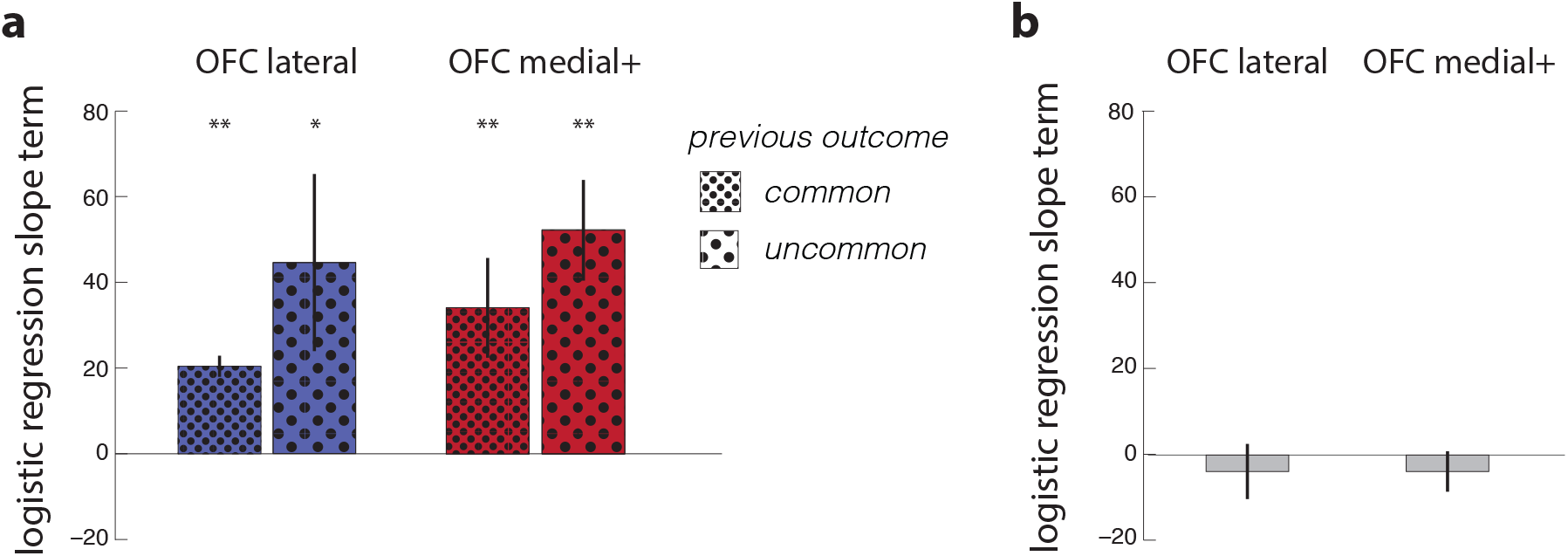
Greater OFC activity at the time of both common and uncommon outcomes was related to higher probability of predicting the common outcome on ensuing trials with the same cue. **(a)** Within-subject, trial-by-trial correlations of OFC activity with learning from recent outcomes, for Cue A and B trials. Plotted is the mean slope term from logistic regression of [probability of predicting the common outcome] on [percent signal change in OFC subregion at the time of outcome, for the most recent trial with the same image cue], fit for each subject separately, and also separately for trials where the previous outcome was the common outcome or where the previous outcome was the uncommon outcome. * p<0.05, ** p<0.01. **(b)** OFC activity at outcomes was not related to the probability of predicting the common outcome on ensuing trials with a different cue. Plotted is the mean slope term from logistic regression of [probability of predicting the common outcome] on [percent signal change in OFC subregion at previous outcome, for the most recent trial with a different image cue], fitted for each subject separately. Bars indicate mean slope terms across subjects ± SEM.

To investigate the possibility that the result in Fig 5a was driven by OFC activity simply indicating an increased level of task engagement or task structure representation, we performed a second, analogous analysis in which we regressed subjects’ predictions on OFC activity at the most recent outcome involving a *different* cue. Note that this regression predicts choice from trials that were, on average, temporally closer than the trials used in the above analysis, since different-cue trials are more common (15.0 ± 0.4 seconds between subsequent different-cue trials, 38.6 ± 1.0 seconds between subsequent same-cue trials). As a result, we might expect the relationship between OFC activity and choice to be even stronger in this analysis. Contrary to this prediction, we found that the regression slope terms were not significantly different from zero, as shown in Fig 5b. Thus OFC activity at the time of an outcome only seemed to affect behavioral predictions on ensuing trials when those trials involved the *same* cue. Together, these analyses suggest that the OFC is involved in updating or reinforcing a learned transition model on a trial-by-trial basis, specifically for the currently observed cue-outcome associations.

## 4 Discussion

Lesion and inactivation studies have shown that the orbitofrontal cortex is particularly important for planning and model-based decision-making. However, prior work implicating OFC in the representation of expected values did not necessarily explain why this area should be important for model-based decision-making in particular, since expected values are generated in both model-based and model-free learning. Here, we have shown that OFC activity at the time of an outcome is related to learning about the transition structure of a task (the tendencies of certain states to lead to other states), which is necessary for accurate planning, shedding new light on the question of why the OFC is critical for model-based decisions.

Using an experimental design that permits constant updating of (probabilistic) transitions between states, we showed that activity in the OFC is correlated with behavioral measures of learning about transition structure. Greater OFC activity at the time of an outcome was positively correlated with the likelihood of optimally predicting the outcome on the next trial with the same image cue, supporting the hypothesized involvement of the OFC in learning of transition structure.

State-transition learning in our experiment was distinct from value-based learning that is thought to be implemented in the dopaminergic system (Jocham et al, 2011; Kravitz et al, 2012), because the trials of interest always led to a predictable number of 2 M&Ms. Our analyses also combined conditions (Cue A and Cue B trials) in which the identities (M&M colors) of the common and uncommon outcomes were reversed, so that any potential differences in value for different M&M colors would cancel out. Therefore, our results positively identify a role for the OFC in learning a non-value-related quantity.

In particular, our results suggest a role for OFC in the activation or reinforcement of an already-learned transition model (and thus the reinforcement of the more optimal prediction), which may be in line with previous work indicating OFC representation of a cognitive map of task space (Wilson et al, 2014; Schuck et al, 2016; Chan et al, 2016; Schuck & Niv, 2019). At the same time, our results are agnostic as to where exactly this model is stored or accessed from – they show only the involvement of OFC in *updating* or *reinforcing* an internal model of transition structure.

What kind of learning might underlie the observed relationships between OFC activity and behavior? Both Hebbian and error-based learning might lead to the pattern of results we observed, though these algorithms would require additional signals to drive the learning process (e.g., from a learned model), in order for increased OFC activity to strengthen the relationship between a cue and the common outcome, rather than the currently observed outcome. Other, yet unknown algorithms, may of course also be possible – in contrast to our increasingly detailed understanding of how world models are used for deploying goal-directed behavior (Daw, 2018), the mechanism for learning world models has not been well-described in the literature as of yet (O’Doherty et al, 2017; Niv, 2019).

Previous work has proposed a state prediction-error algorithm for learning state transitions, analogous to learning about state values from reward prediction errors observed in dopaminergic neurons. A state prediction error would signal surprise-related information at the time of an unexpected state (regardless of the state’s value), and could be used to adjust internal estimates of transition probabilities by increasing the estimated probability of transitioning to the just-observed state. However, our results did not straightforwardly affirm the hypothesis that the OFC supports state-transition learning by representing a state prediction error signal. At the least, we did not find evidence for encoding of this error signal in the OFC’s univariate response to outcomes, as we did not observe differences in OFC activity for common vs. uncommon outcomes (corresponding to small vs. large state prediction errors). Further, univariate OFC activity at the time of an outcome was not correlated with greater subsequent expectations of that particular outcome. Instead, OFC activity was related to greater subsequent expectation of the *more common* outcome (i.e. more optimal prediction by subjects), regardless of whether the OFC activity was measured at the time of a common or uncommon outcome.

Previous works have found mixed evidence of state prediction errors being represented in the OFC. Howard and Kahnt (2018) and Suarez et al. (2019) discovered evidence of state prediction errors in the midbrain, OFC, piriform cortex, amygdala, lateral prefrontal cortex, and posterior parietal cortex. Boorman et al. (2016) found correlations of state prediction errors with univariate activity in lateral OFC, but only when signed positively or negatively according to whether the update increased or decreased the odds of a preferred outcome. However, Gläscher et al. (2010) tested for univariate correlations with the (unsigned) magnitude of an inferred state prediction error signal, and implicated the dorsolateral prefrontal cortex and intraparietal sulcus (but not the OFC) in their analysis.

It is possible that the discrepancy between our findings and previous work showing state prediction errors in OFC is due to the specific differences between our experiment and previous work – differences that perhaps highlight the specific circumstances required for observation of state prediction errors in the OFC. First, state transition probabilities in the current experiment (and in Glascher et al, 2010) were stationary (though stochastic), whereas the aforementioned previous works used non-stationary distributions with sudden reversals in cue-outcome contingencies. Recent work has indicated that humans and animals interpret large changes in the environment differently from small surprises, with large surprises taken as indicating a new underlying “latent state” of the environment (Gershman et al, 2010; Collins and Frank, 2013; Dunsmoor et al, 2015, Gershman et al., 2017). The process of latent state inference and representation has been associated with both the hippocampus and the OFC (Schuck & Niv, 2019; Niv 2019).

A second difference is that our main analyses averaged over any differences in value for different outcomes by aggregating across Cue A and Cue B, in order to disentangle reward prediction errors and state prediction errors. Glascher et al also analyzed state prediction errors only in the absence of rewards, and did not find evidence for state prediction errors in OFC. Indeed, for the trial type where we purposefully did not take these measures to disentangle state and reward prediction errors (Cue D trials, where the image cues led probabilistically to varying numbers—rather than colors—of M&Ms), we did find evidence of univariate differences in activation of lateral OFC for uncommon vs. common (i.e. high vs. low value) outcomes. This is reminiscent of the results from Boorman et al (2016), who found correlations with state prediction errors in lateral OFC only when the inferred errors were signed positively or negatively according to the change in value of a state transition. Howard & Kahnt (2018) also showed that responses to (unsigned) identity and positive value prediction errors were correlated across participants, at least in the midbrain. These results suggest some kind of shared neural response between state and reward prediction errors in these studies.

Previous work in rats that implicated the OFC in learning about transition structure, concentrated on the lateral OFC (McDannald et al, 2010), and fMRI work in humans also specifically implicated the lateral OFC in this type of process (Boorman et al, 2016; Howard and Kahnt, 2018; Suarez et al, 2019). We tested our hypotheses in the entirety of the OFC, using a previously determined functional connectivity-based parcellation of OFC into medial and lateral subregions (Kahnt et al, 2012). Medial and lateral OFC showed very similar results across all our analyses. Of course, this does not rule out the possibility that there may exist a different parcellation of OFC that would lead to differing results across subregions. We note also that the homology of OFC between rodents and humans is currently unclear, and OFC subdivisions are particularly complex given observed considerable anatomical variability within individuals (Wallis et al, 2011; Chiavaras and Petrides, 2000). We should also take care in interpreting the negative BOLD response in OFC – this negative BOLD response has been previously observed (e.g. Boorman et al, 2009), but is not yet fully understood.

It is important to note that our secondary analyses did provide further support for two other mainstream theories of OFC function, in addition to the theory of a role in learning transition structure. First, we found that we could decode representation of outcome identity in OFC using multivariate pattern analysis (MVPA) of BOLD activity at the times of the outcomes, consistent with a recent theory implicating OFC in representation of the current state (Wilson et al, 2014), for which evidence is increasingly amassing (e.g. Klein-Flügge et al, 2013; Bradfield et al, 2015; Chan et al., 2016; Schuck et al, 2016; Nogueira et al, 2017; Howard et al, 2020; Zhou et al, 2020). Second, we found evidence for value sensitivity in univariate BOLD responses in lateral OFC in a separate task condition (Cue D), in which the number (but not color) of M&Ms was unpredictable, consistent with previous work demonstrating that OFC represents the value of rewards (Gottfried et al, 2003; Padoa-Schioppa and Assad, 2006; Hampton et al, 2006; Fellows, 2007; Hare et al, 2008; Wallis and Kennerley, 2011; Monosov and Hikosaka, 2012; though note that value might be construed as just one feature in representation of the current state; Lopatina et al, 2015).

In conclusion, the present results provide support for an emerging understanding of the relationship between the OFC and acquisition of state-to-state transition structure. Our findings may suggest a role for OFC in reactivation and reinforcement of an already learned state-transition model, relating to proposals that the OFC stores such a model (Wilson et al, 2014). Our findings also build upon previous work showing that rats with OFC lesions show impaired learning about changes in state transitions (McDannald et al, 2011), and that surprise signals in human OFC are related to changes in hippocampal representations of state transitions (Boorman et al., 2016). Importantly, while our results are not straightforwardly aligned with a state prediction error hypothesis, at least in this learning regime, they do still indicate OFC’s involvement in learning about transition structure. Our findings may therefore serve to constrain future models of the particular learning algorithms that may underlie learning about transition structure, facilitating a fuller understanding of the involvement of OFC in learning and model-based decision making.

